# A frosty genetic screen unmasks a major regulatory role for SHORT VEGETATIVE PHASE of flowering in response to a cold snap

**DOI:** 10.1101/2024.03.20.585907

**Authors:** Ashleigh Edwards, Hans Thordal-Christensen, Stephan Wenkel

## Abstract

The control of flowering in plants is intricately governed by a combination of internal and environmental signals, with temperature playing a critical role. Thus, *Arabidopsis thaliana* plants display temperature-dependent variations in flowering time. As unexpected periods of cold temperatures can occur at any time, plants have evolved mechanisms to detect such cold snaps and to respond by delaying flowering. Plants are more tolerant to cold temperatures in the vegetative stage, while flowers are more sensitive and have reduced reproductive success due to damage to floral structures and gametes. At the molecular level, delayed flowering can be caused by repressing the *FLOWERING LOCUS T* (*FT*) gene, and several MADS box transcription factors have been shown to repress *FT* expression in response to cold and in this way prevent flowering. Here, we employed a forward genetic screen aimed at understanding the effect of a cold snap on the transition to flowering. We germinated a population of *A. thaliana* EMS M2 plants at 20°C and then gradually lowered the temperature to 10°C and selected early flowering mutants. Using whole-genome sequencing, we identified seven mutant alleles of the *SHORT VEGETATIVE PHASE* (*SVP*) gene. This finding establishes a central role for *SVP* in repressing flowering in response to a cold snap and provides novel alleles, several of which affect splice junctions. Our research thus presents valuable insights into the nuanced molecular mechanisms governing temperature-responsive flowering in Arabidopsis and sheds light on the dynamic interplay between *SVP* and environmental cues.

## Introduction

Based on the third law of thermodynamics, an increase in temperature results in a corresponding increase in entropy. Biological processes typically adhere to the laws of physics, and the movement of molecules or enzymatic reactions can vary as a function of temperature. Complex physiological responses, comprising numerous factors, are no exception. However, in addition to slowing down the movement of molecules or reducing the fluidity of membranes, biological systems have evolved other forms of negative regulation in response to low temperatures. The process of flowering is a complex physiological response regulated by an intricate combination of endogenous and environmental signals. Temperature significantly affects flowering. Some vernalization-dependent accessions of *Arabidopsis thaliana* require winter cold to initiate flowering in the next spring. Other accessions, including the widely studied Col-0 accession, are vernalization-independent mutants that flower rapidly without cold-treatment. However, they show changes in flowering time when exposed to ambient cold (Kim et al., 2009). Under laboratory conditions, Arabidopsis is typically grown at around 20°C, causing Col-0 plants to flower early when photoperiods are long (e.g. 16 hours day, 8 hours night). Under short days (e.g. 8 hours day, 16 hours night), flowering is severely delayed because the photoperiodic flowering time pathway is not active (Andres and Coupland, 2012). Elevated temperatures (e.g. 28°C) cause very early flowering under long days and a significant acceleration of flowering under short days, which is attributed to the activation of the florigen *FLOWERING LOCUS T* (*FT*) by PHYTOCHROME INTERACTING FACTOR4 (PIF4) (Kumar et al., 2012).

The Arabidopsis Col-0 accession also shows variation in flowering in response to ambient cold, and flowering is strongly delayed in cold temperatures, even when days are long (Blázquez et al., 2003). Several transcription factors from the MADS-box family that are closely related to the master vernalization regulator FLOWERING LOCS C (FLC) have been identified as significant regulators of flowering in response to ambient cold. One of these transcription factors is SHORT VEGETATIVE PHASE (SVP), which inhibits flowering at low temperatures, preventing plants from flowering under suboptimal conditions. Plants carrying mutations in the *SVP* gene exhibited an early flowering phenotype that was more pronounced when plants were grown at 16°C compared to 23°C (Lee et al., 2007). At the molecular level, SVP acts as a repressor of *FT* expression and overexpression results in late flowering plants (Lee et al., 2007). The interaction between SVP and FLC causes the repression of flowering, and both transcription factors have interdependent and overlapping functions and distinct tissue specificities (Mateos et al., 2015). Moreover, the SVP-FLC complex acts as a negative regulator on many activators of flowering time, as well as genes that encode enzymes responsible for the production of the hormone gibberellic acid (Andrés et al., 2014; Mateos et al., 2015).

Another gene belonging to the MADS-box family, *FLOWERING LOCUS M* (*FLM*), is also involved in the regulation of flowering in response to temperature (He et al., 2004). However, the *FLM* gene makes several splice isoforms, with *FLM-β* and *FLM-δ* emerging as the major ones in the Col-0 accession, and these isoforms exert opposing influences on the regulation of flowering time (Lee et al., 2013; Posé et al., 2013). *FLM-β* operates as a floral repressor resulting in delayed flowering, whereas *FLM-δ* induces the opposite effect by promoting early flowering. The interplay of these isoforms depends on the ambient temperature. *FLM-β* is favored in lower temperatures, which suppress flowering. On the other hand, *FLM-δ* is more abundant in higher temperatures, alleviating repression, and promoting early flowering (Lee et al., 2013; Posé et al., 2013). Furthermore, the FLM-β isoform facilitates the translocation of SVP into the nucleus at low temperatures, whereas at elevated temperatures, the FLM-δ isoform interacts with SVP and retains it in the cytoplasm (Jin et al., 2022). This dual feedback-control mechanism guarantees that flowering is inhibited in the cold and accelerated in high temperatures.

Here, the effect of a cold snap on the transition to flowering was investigated using a forward genetic screen. The objective was to identify regulators that repress flowering in response to cold. We reasoned that it would be advantageous to know such regulators because, with climate change, such mutants could adapt more quickly to warmer winters, ensuring timely flowering and more predictable and stable yields. In regions with short growing periods, such as the subarctic North, plants that can endure cold spells without delaying flowering can enhance plant productivity.

We therefore germinated an Arabidopsis mutant population at 20°C under long day conditions and, after three days, lowered the temperature by 2°C each day to 10°C for screening purposes. As a result, a total of 12 mutants displaying early flowering under these conditions were isolated with no other discernible side effects.

Experiments investigating the flowering time of the mutants and wild type were conducted under short- and long-day conditions at 22°C, showing a mild early flowering phenotype of the mutants under long day conditions and a stronger effect under short day conditions. Following back-crossing of five *frosty* mutants to Col-0 and self-pollination, segregating offspring were grown in cold long day conditions and genomic DNA was extracted from up to 30 early flowering individuals per mutant line. Whole-genome sequencing of pooled DNA showed that all mutants carried mutations in the *SVP* gene. Therefore, our study has identified *SVP* as a significant suppressor of flowering in response to cold snap. In total, we present seven new alleles, a subset of which affects splice junctions.

## Materials and Methods

### Plant material and growth conditions

Plants were accession Col-0 and *frosty* mutants were identified from EMS mutagenized populations containing the *eds1*-*2* mutation (Hans Thordal-Christensen, *unpublished*). To assay for early flowering, seeds were first stratified on soil for three days before being placed in one of the following three growth regimes: long-day (16-hr day, 8-hr night) with light intensity of 150 μmol·m^-2^·s^-1^, temperature of 21°C during day and 19°C during night, and humidity of 60% during day and 65% during night; short-day (8-hr day, 16hr night) with light intensity, temperature, and humidity identical to the long-day regime; or cold long-day (16-hr day, 8-hr night) with light intensity identical to the long-day regime, but only the initial three days with a constant temperature of 21°C after which the temperature was decreased by 2°C per day to reach a constant temperature of 10°C, with a constant humidity of 60%. Flowering time was determined by the number of rosette leaves at bolting.

### Genotyping of *eds1-2*

Genomic DNA was extracted from individual F1 plants following the Edward’s protocol for DNA extraction (Edwards et al., 1991) and 2 µl was used for PCR amplification of *EDS1* using forward oligo 5’-ACACAAGGGTGATGCGAGACA-3’ with reverse oligo 5’-GTGGAAACCAAATTTGACATTAG-3’. For the amplification we used Taq polymerase with an annealing temperature of 59°C and extension time of one minute.

### Whole genome sequencing library preparation

To prepare DNA for whole genome sequencing, single leaf discs were collected and pooled from between 20-30 early flowering individuals of *frosty1, frosty6, frosty7, frosty8, and frosty10*. At the same time, single leaf discs were collected and pooled from 20 individual randomly selected *eds1-2*/+ plants. Genomic DNA was extracted using the DNeasy Plant Pro Kit (Qiagen). For the DNA library preparations, we used only 0.2 μg DNA per sample. First, we sonicated the genomic DNA sample to get fragments of 350 bp size. Then, we performed end-polishing, A-tailing and ligated the fragments with the full-length adapter specified for Illumina sequencing.

Additional size selection and PCR amplification followed. Lastly, we purified the PCR products using the AMPure XP system in Beverly, USA. The library’s quality was tested with Agilent 5400 in the United States and then measured using QPCR (1.5 nM). The libraries that passed the evaluation were brought together and sequenced on Illumina platforms utilizing the PE150 approach, considering the concentration of the library and the needed amount of data.

### Mapping and SNP calling

Quality-checked sequencing data was aligned to the reference genome through the use of BWA software (Li and Durbin, 2009). This helped obtain initial mapping outcomes saved in the BAM format by setting the parameter: mem -t 4 -k 32 -M. To enhance the accuracy of the results, redundancies were cleared using the SAMtools (Li et al., 2009) and Picard tools (http://broadinstitute.github.io/picard/).

GATK (DePristo et al., 2011) was used to detect the initial SNP and InDel sets, using settings such as ’– gcpHMM 10 -stand_emit_conf 10 -stand_call_conf 30’. Subsequently, we filtered these groups according to the following standards: SNP: QD < 2.0, FS > 60.0, MQ < 30.0, HaplotypeScore > 13.0 MappingQualityRankSum < -12.5, ReadPosRankSum < -8.0 INDEL: QD < 2.0, FS > 200.0, ReadPosRankSum < -20.0 ANNOVAR (Wang et al., 2010) was used to annotate variants, with gene and region annotations based on the known genes from UCSC.

### Detection of *SVP* splice isoforms

Total RNA was extracted from two rosette leaves of individual M2 plants following protocol A of the Spectrum™ Total RNA Kit and genomic DNA then removed using the Thermo Scientific™ RapidOut DNA Removal Kit. 650 ng of RNA was used for cDNA synthesis which was done using the iScript™ cDNA Synthesis Kit. To amplify the entire coding sequence of *SVP*, we used forward oligo 5’-GCTCTCTCTCTTGCTTCTAG-3’ and reverse oligo 5’-CCTCACTCCATCAACTTCTT-3’. PCR amplification was done using Phusion® High Fidelity DNA Polymerase (NEB) with an annealing temperature of 60°C and an extension time of 30 seconds. PCR products were visualized on a 1% agarose gel and purified using the E.Z.N.A® Cycle Pure Kit. Purified DNA was sequenced using Azenta Life Sciences Genewiz Sanger Sequencing services.

## Results

### Identification of *frosty* mutants that flower rapidly in response to a cold snap

The Col-0 ecotype of Arabidopsis exhibits substantial suppression of flowering at reduced ambient temperatures. A genetic screen was conducted to identify regulators that suppress flowering in response to a cold snap. For this study, we utilized the Col-0 *eds1-2* mutant (Bartsch et al., 2006), which is highly susceptible to pathogens, and mutagenized it with EMS to induce random point mutations (Hans Thordal-Christensen, *unpublished*). Prior to mutagenesis, extensive backcrossing of *eds1-2* into the Col-0 from L*er* (Bartsch et al., 2006) allowed us to use Col-0 for backcrossing with potential *frosty* mutants and provided us with a genetic marker to validate the success of these genetic crosses.

To identify suppressors of flowering in cold snap conditions, we reasoned that we needed to ensure that the plants germinated uniformly and then rapidly adapted to the colder ambient temperatures. To achieve this goal, we have developed a cold snap growth regime (**Fig. 1A**). Before germination, seeds were sown on soil and exposed to 4°C in darkness for three days to break dormancy. Then, the seeds were permitted to germinate under long-day conditions at a temperature of 22°C for three full days, followed by a gradual decrease in temperature of 2°C per day until 10°C was reached. We assessed the flowering behavior of around 200,000 plants from 20 mutagenized seed batches under these cold long-day conditions and identified early flowering mutants from six different seed pools. Some of these mutants showed extremely early flowering, reduced plant size and reduced fertility. As these pleiotropic mutants may affect genes that encode factors involved in diverse processes, our focus was on 12 *frosty* mutants that showed only early flowering without any other significant side effects.

**Figure 1.**
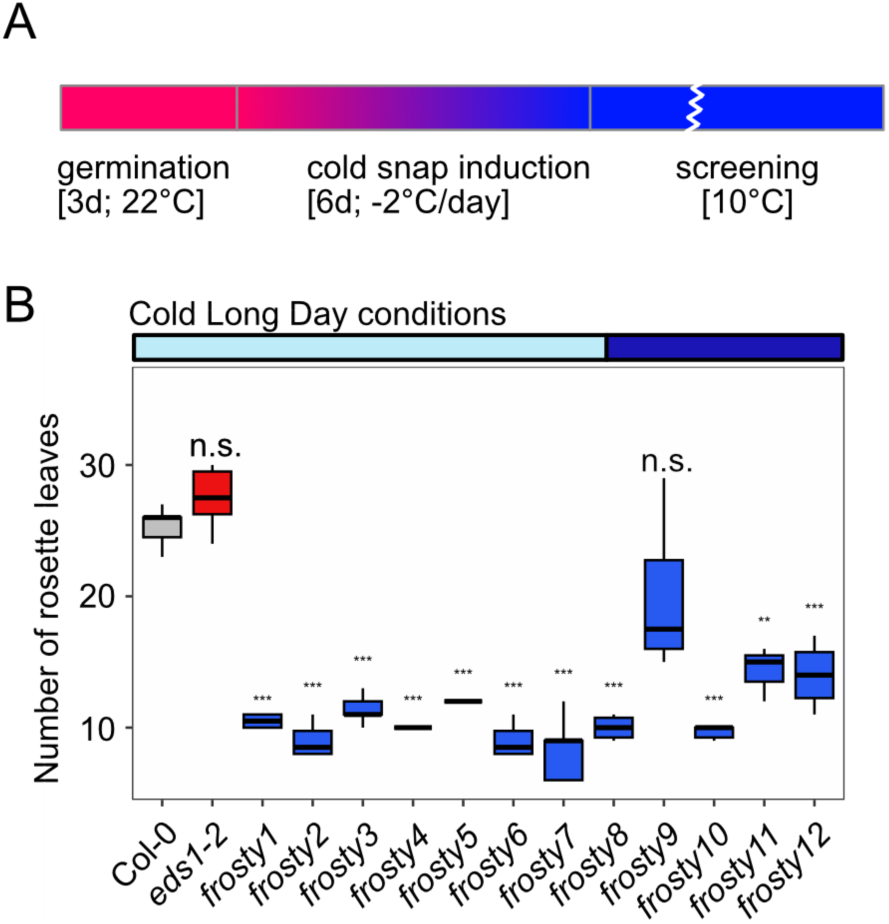
Identification of *frosty* mutants. **(A)** Schematic overview of the flowering time experiment under cold conditions. After a short 3-day germination phase at 22°C, the plants were exposed to a cold snap by gradually cooling the growth chamber at 2°C per day until the screening conditions reached 10°C. Plants were cultivated at 10°C until the transition to flowering and subsequent seed set. **(B)** Quantifying flowering time in cold long days (16h light, 8h dark, 10°C) was based on counting the number of rosette leaves at the transition to flowering. The central line on the box plots denotes the median, while the box limits indicate the 25th and 75th percentiles. The whiskers extend 1.5 times the interquartile range from the 25th and 75th percentiles. Asterisks indicate significance levels between values that were determined through a two-sample T-Test. For all plots, **P ≤ 0.01, ***P ≤ 0.001, and n.s. denotes insignificance.

To confirm that these 12 *frosty* mutants were homozygous and early-flowering, we collected their seeds and grew their offspring under identical conditions. We compared the flowering time of these mutants with the Col-0 wild type and the *eds1-2* mutant plants which represented the genetic background of all the *frosty* mutants. With the exception of the *frosty9* mutant plants, all mutants displayed consistent early flowering in cold long-day conditions (**Fig. 1B**). Given the uniformity of the flowering responses of the other mutant plants, it appeared that they were homozygous for mutations in potential *FROSTY* genes. The *eds1-2* mutant plants exhibited the same flowering time as the Col-0 wild type plants. Consequently, our study has identified several mutants that exhibit the loss of floral repression in response to a cold snap growth regime.

### Flowering time analysis of *frosty* mutants grown under short- and long-day conditions at warm temperatures

Daylength is a major determinant of the flowering time of Arabidopsis Col-0 plants grown under standard growth conditions (here 22°C). To determine whether the *frosty* mutants exhibit any changes when cultivated under typical long- and short-day conditions, we cultivated the corresponding mutants in these conditions.

When grown under warm long day conditions, all frosty mutants, apart from *frosty7*, *frosty9* and *frosty11*, were found to be showing significant early flowering behaviors (**Fig. 2A**). As in the cold long day conditions, the *eds1-2* mutant flowered indistinguishably from the wild type. In short-day conditions all *frosty* mutant plants flowered early compared to the control plants (**Fig.2B**). We did observe a trend for *eds1-2* mutants to flower somewhat earlier than the Col-0 wild type, albeit not significantly. The weakest early flowering phenotype, with substantial variability, was observed in the *frosty9* mutant plants, which may indicate a multi-gene effect in this line. Additionally, *frosty3* and *frosty9*, displayed reduced fertility and were therefore removed from further analysis. The findings suggest that the causal mutations affect negative regulatory factors that suppress flowering. This is supported by our earlier observations that *frosty* mutants rapidly flower in cold long day conditions.

**Figure 2.**
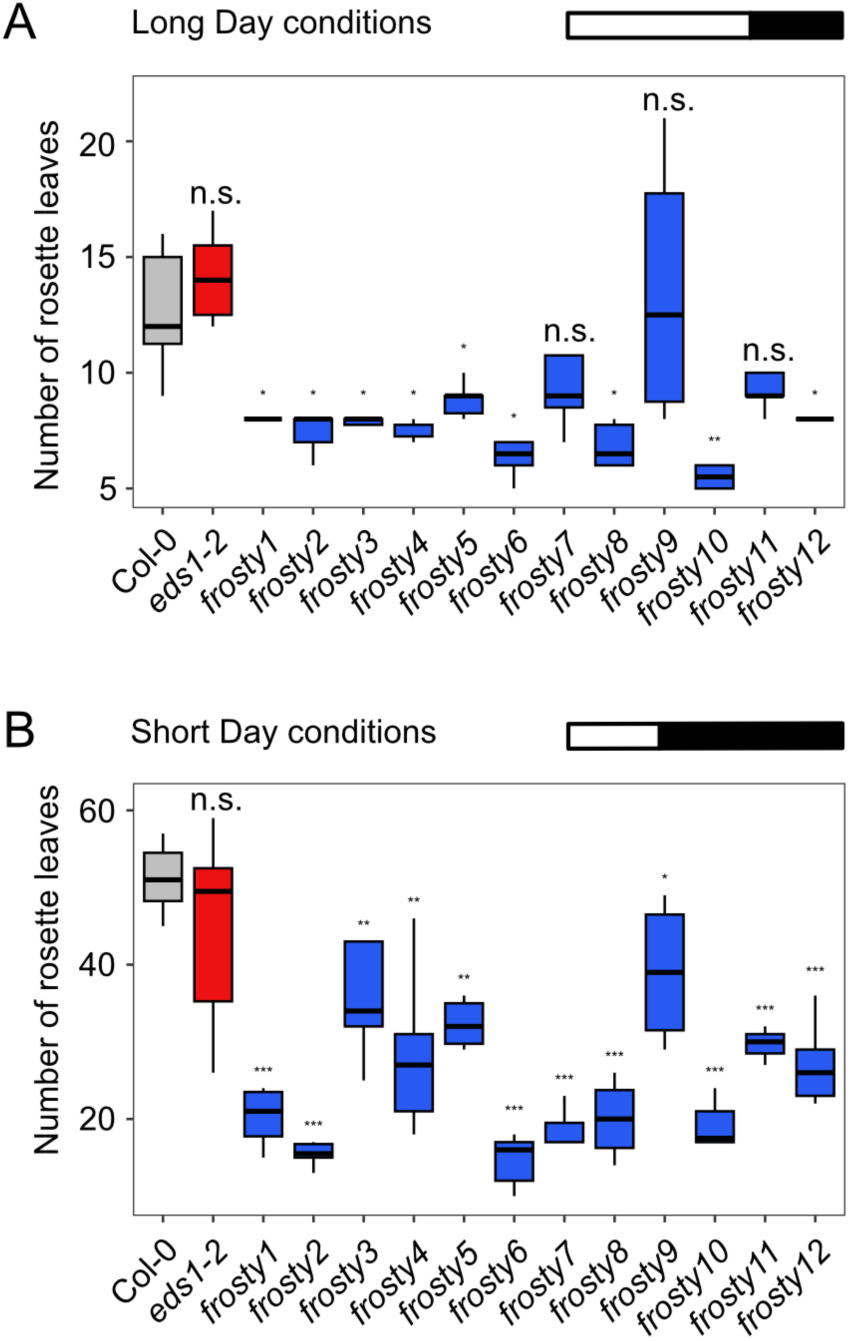
Flowering time analysis of *frosty* mutants grown in warm short- and long-day conditions. Quantification of flowering time in **(A)** long- and **(B)** short day conditions by counting the number of rosette leaves produced at the transition to flowering stage. The median is represented by the central line in the box plots, with the box boundaries indicating the 25th and 75th percentiles. The whiskers extend 1.5 times the interquartile range from the 25th and 75th percentiles. Any significance levels observed between values are denoted by asterisks within the plots: *P ≤ 0.05, **P ≤ 0.01, ***P ≤ 0.001, and n.s. indicates insignificance.

### Genetic analysis aimed at revealing the molecular basis of the *frosty* mutants

EMS mutagenesis induces thousands of random mutations in the genome. As a result, the identification of plants exhibiting traits of interest may be attributed to multigenic effects. To identify the causal genetic mutations and establish the monogenic nature of the trait, it is necessary to outcross the mutant plants and analyze the segregation patterns in the subsequent self-pollinated progeny. As EMS mutagenesis was conducted on *eds1-2* mutant plants and all *frosty* mutants are also homozygous *eds1-2* mutants, the *EDS1* gene was employed as a marker for successful backcrossing (**Fig.3**).

**Figure 3.**
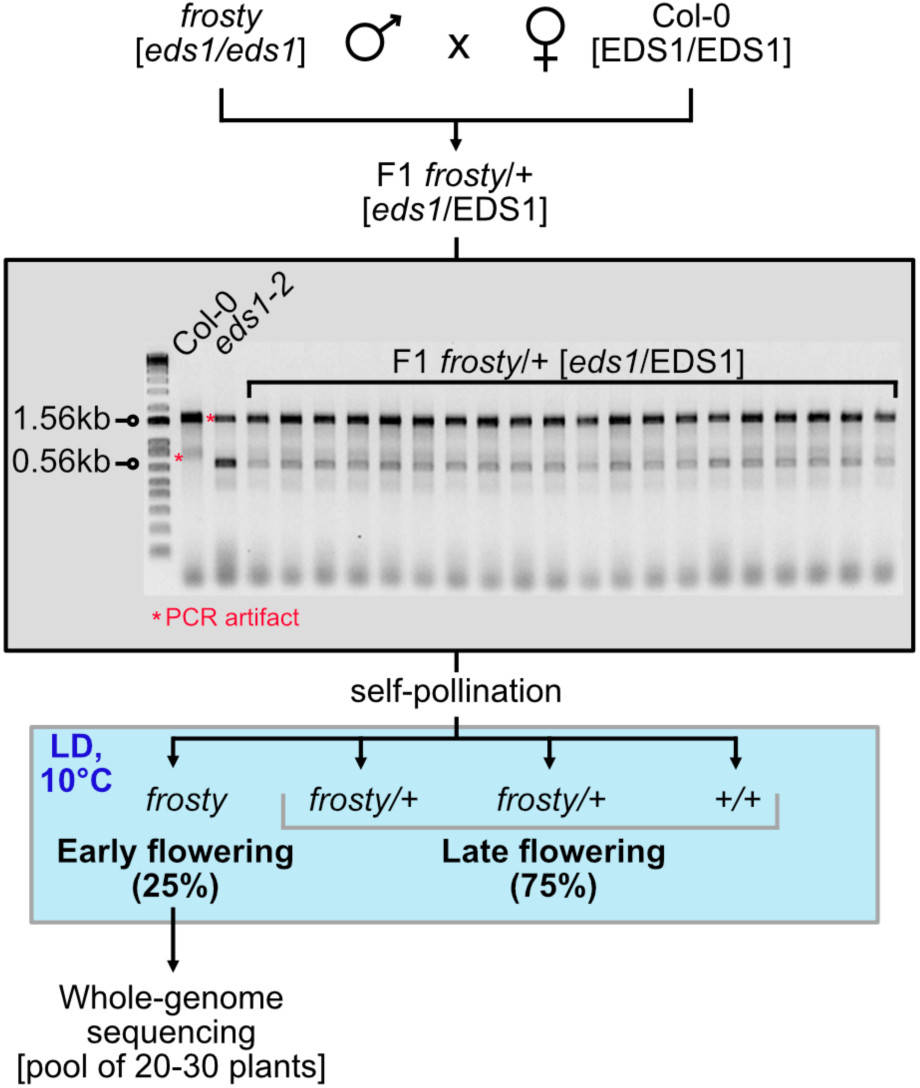
Graphic representation of the genetic analysis of *frosty* mutants. To conduct the analysis, Col-0 wild type plants were fertilised with *frosty* mutant pollen. The resulting F1 plants were PCR tested using primers that amplify in the *EDS1* locus. In Col-0 wildtype, a 1.56kb fragment is produced, whilst in the *eds1-2* mutant, a 0.56kb fragment is produced. However, the *eds1-2* mutant also generates a spurious PCR product at 1.5kb, which prevents the use of Col-0 pollen on *eds1-2* mutants. The resulting F1 genotyping unambiguously shows that all Col-0 wild-type plants have been successfully pollinated by frosty *eds1-2* mutant pollen. Afterwards, heterozygote plants were self-pollinated and the progeny was grown in cold long days. The genetic material of *frosty* mutants, exhibiting a clear Mendelian segregation, was collected for whole-genome sequencing.

In our genetic experiments, we cross-pollinated Col-0 wild-type plants with pollen from the different frosty mutants. Due to polymorphisms in the *EDS1* gene, we could confirm the heterozygous nature of the resulting offspring using PCR (**Fig. 3**). These heterozygous F1 *frosty* mutant plants, all displaying wild-type flowering time (data not shown), were self-pollinated and their individual offspring were grown in cold long-day conditions to rediscover the fraction of homozygous early-flowering plants. In summary, our genetic analysis revealed that some *frosty* mutant plants (see below) may be the result of a single mutation in a factor that acts as a floral repressor, which segregates recessively in the respective mutant populations.

### Whole genome-sequencing and SNP analysis to identify the molecular nature of the *frosty* mutants

To find the mutations responsible for early flowering under cold long-day conditions, we grew F_2_ seeds from self-pollinated plants (**Fig.3**) under the conditions described above (**Fig.1A**). As experimental controls for flowering, we again included the Col-0 wild type, the *eds1-2* mutant and an F2 generation resulting from a cross between Col-0 and *eds1-2*. The latter was included as a reference for the subsequent whole genome sequencing analysis.

Counting the number of rosette leaves to quantify the onset of flowering revealed a clear fraction of early flowering plants in the *frosty1*, *frosty6*, *frosty7*, *frosty8* and *frosty10* mutant pools (**Fig.4A**). Determining which plants had early flowering characteristics became more difficult in the remaining *frosty* mutant pools. These had a mixture of plants with characteristics similar to control plants and were therefore excluded from whole genome sequencing. Our subsequent whole genome sequencing experiments focused therefore on *frosty1*, *frosty6*, *frosty7*, *frosty8* and *frosty10* mutant plants, which showed apparent Mendelian segregation, suggesting a monogenic trait (**Fig.4A,B**).

**Figure 4.**
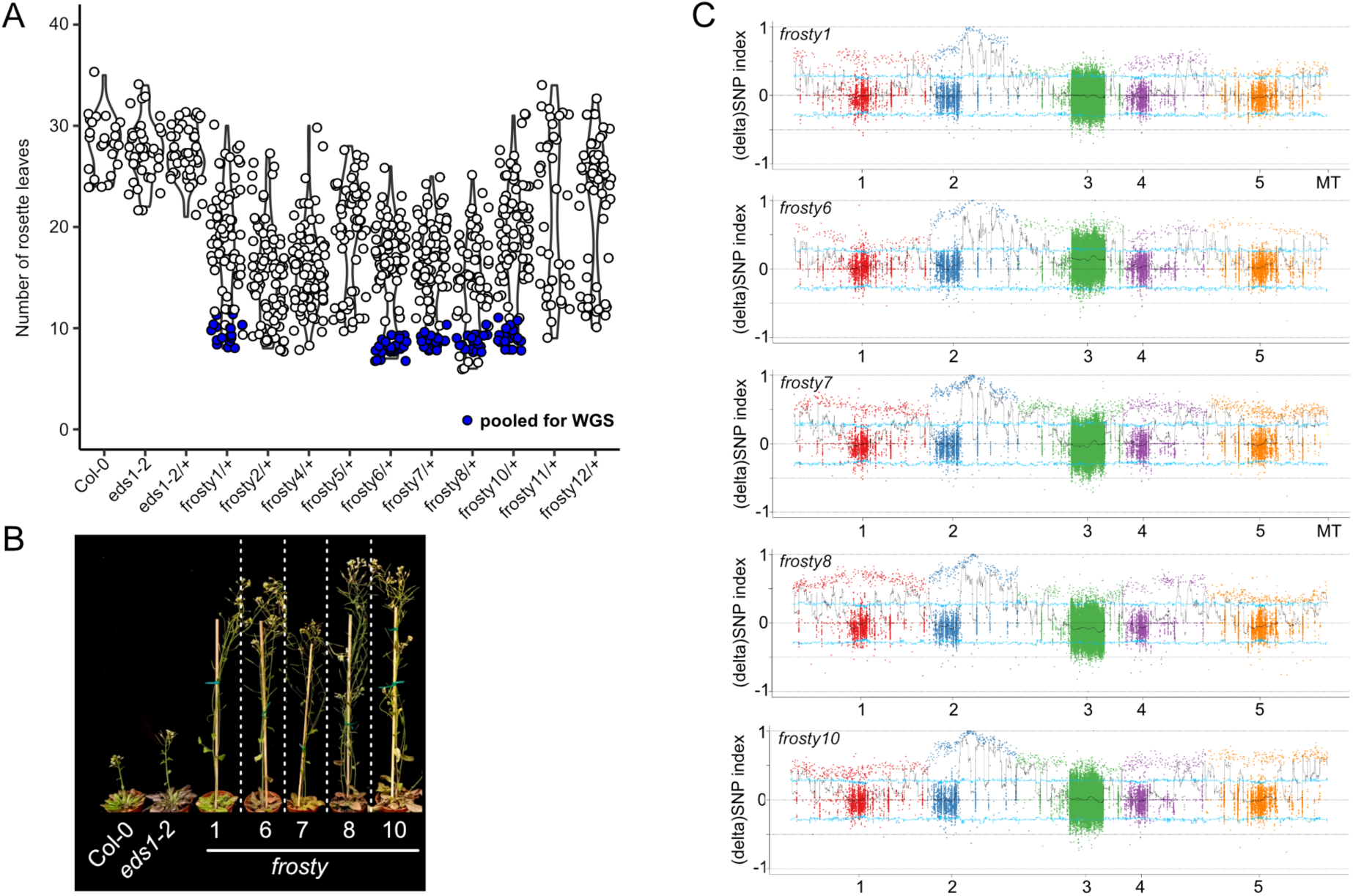
Molecular cloning of *frosty* mutants. **(A)** Flowering time analysis of self-fertilized progeny of individual *frosty/+ eds1-2/+* mutant plants. The dark-blue dots represent individual plants whose genomic DNA was isolated for pooled whole-genome DNA sequencing. **(B)** Pictures of representative plants selected for whole-genome re-sequencing. **(C)** ΔSNP indexes of mutant pools indicating that all mutant pools have a SNP-frequency close to 1 on chromosome 2.

Leaves were harvested from 20-30 F_2_ individuals per *frosty* mutant pool, and the genomic DNA was fragmented to approximately 350bp. After ligation of the Illumina adapters, the libraries were deep sequenced using 150bp paired-end sequencing. On average, 104 million reads were obtained per sample equating to 15.5Gb of cleaned data or 100X the size of the entire genome, which corresponds to 99.99% coverage.

After mapping all reads to the genome, we compared the SNPs identified in *eds1-2/+* with the SNPs identified in the individual *frosty* pools. By plotting the differences in SNP frequency between the individual *frosty* mutant pools and *eds1-2/+*, we observed a notable increase in SNP allele frequencies on chromosome 2 (**Fig. 4C**).

Further examination of the SNP peak revealed that all sequenced *frosty* mutants had at least one mutation in the *SHORT VEGETATIVE PHASE* (*SVP*) locus *AT2G22540* in 100% of the reads and SVP is known to suppress flowering at low temperatures (Lee et al., 2007). In the *frosty1* mutant, we identified a second homozygous single nucleotide polymorphism that results in a premature stop codon in the gene for *DIACYLGLYCEROL KINASE 5* (*DGK5*; *AT2G20900*). Similarly, in *frosty8*, a premature stop codon was found in a gene encoding a cysteine/histidine-rich C1 domain family protein (*AT2G21850*). Additionally, *frosty10* carried a second mutation site in the gene for *MITOCHONDRIAL EDITING FACTOR 21* (*MEF21*; *AT2G20540*).

It is worth noting that all of the second site mutations are found in close vicinity of *SVP* and are therefore likely to be the result of genetic linkage and not causal for the flowering phenotype. This hypothesis is supported by the fact that none of the corresponding genes has a known role in flowering regulation. Further support for identification of *SVP* as a *FROSTY* gene came from finding of two additional EMS mutant *svp* alleles in *frosty2* and *frosty4* by direct Sanger sequencing. We therefore conclude that the identified mutations in the *SVP* locus are causal for the observed flowering promotion.

### Identification of novel SVP mutant alleles

Database and literature searches of the *SVP* locus uncovered several loss-of-function mutations that had been previously identified. Various T-DNA insertion lines have been analyzed, all located within non-coding regions (5’UTR and introns) (**Fig. 5A** and **Table 1**). However, because of the T-DNA’s considerable size, all T-DNA mutants seem to represent complete knock-outs of *SVP* (Hartmann et al., 2000; Lee et al., 2007; Bechtold et al., 2016). A series of frameshift mutants (*svp41*, *svp42* and *svp43*) in the first exon were isolated via transposon mutagenesis, leading to premature termination of transcripts and representing null alleles. Additionally, mutant alleles were identified in genetic screens utilizing EMS as a mutagen. These mutants typically exhibit an early flowering phenotype. Interestingly, numerous EMS-induced mutations affect splice junctions, indicating that SVP is susceptible to mis-splicing and more robust to missense mutations in the coding sequence.

**Figure 5.**
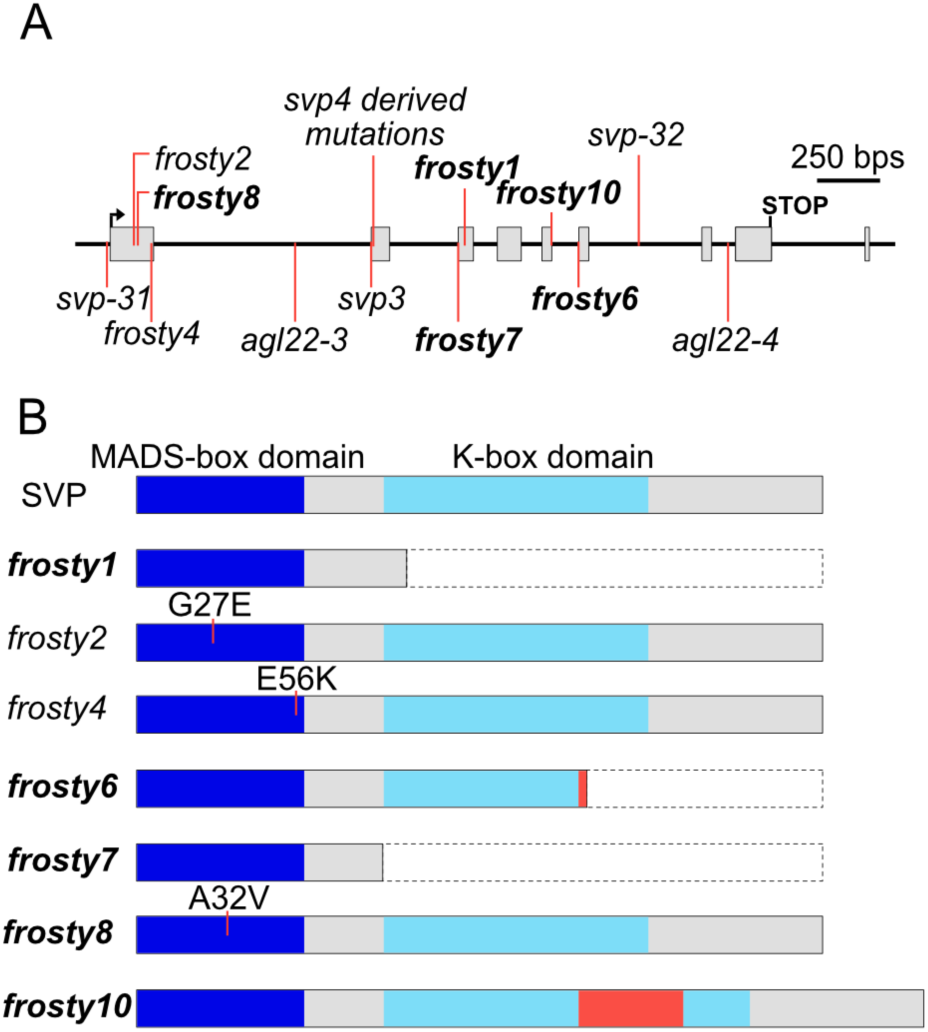
Overview of mutations in the *SVP* gene. **(A)** The genomic structure of *SVP* is represented by grey boxes which indicate exons, and red lines which indicate the positions of mutations. The SVP protein structure is depicted in **(B)**, with the MADS-box domain shown as a dark blue box and the K-box domain shown as a light blue box. Other protein sequences are represented by grey boxes. Red boxes indicate novel protein sequences resulting from alternative splicing, while the dotted white boxes represent sequences that have been lost due to the creation of premature stop codons. Mutants highlighted in bold were identified through whole-genome re-sequencing.

**Table 1.** Overview of alleles of SVP so far characterised.

| Allele | Type | Chromosomal position | Effect | Phenotypes besides early flowering | Background | Reference |
| --- | --- | --- | --- | --- | --- | --- |
| **svp-4** | T-DNA insertion | 9,581,517^9,581,518 | Knock-out |  | Col-0 | Hartmann et al., 2000 |
| **svp-41 (svp-4 derived)** | 2-bp deletion | 9,581,516-9,581,517 | Frameshift, truncation | reduced response to vernalization in long-day |  |  |
| **svp-42 (svp-4 derived)** | 4-bp insertion | 9,581,517^9,581,518 | Frameshift, truncation |  |  |  |
| **svp-43 (svp-4 derived)** | 10-bp deletion | 9,581,516-9,581,525 | Frameshift, truncation |  |  |  |
| **svp-1** | T-DNA insertion | Unknown | Loss-of-function |  | Ws | Scortecci et al., 2003 |
| **svp-31 (SALK_026551)** | T-DNA insertion | 9,580,406^9,580,407 | Knock-out |  | Col-0 | Lee et al., 2007 |
| **svp-32 (SALK_072930)** | T-DNA insertion | 9,582,644^9,582,645 | Knock-out |  |  |  |
| **svp-3** | EMS mutation | 9,581,505 | Splice site mutation |  | Ler | Fujiwara et al., 2008 |
| **agl22-3 (SALK_141674)** | T-DNA insertion | 9,581,201^9,581,202 | Knock-out | Increased rosette size, increased tolerance to drought stress | Col-0 | Bechtold et al., 2016 |
| **agl22-4 (SAIL_853_C08)** | T-DNA insertion | 9,582,996^9,582,997 | Knock-out |  |  |  |
| **frosty1** | EMS mutation | 9,581,895 | Nonsense (R96X) |  |  | Here |
| **frosty2** | EMS mutation | 9,580,496 | Missense (G37E) |  |  |  |
| **frosty4** | EMS mutation | 9,580,582 | Missense (E56K) |  |  |  |
| **frosty6** | EMS mutation | 9,582,388 | Splice site mutation |  |  |  |
| **frosty7** | EMS mutation | 9,581,870 | Splice site mutation |  |  |  |
| **frosty8** | EMS mutation | 9,580,511 | Missense (A32V) |  |  |  |
| **frosty10** | EMS mutation | 9,582,278 | Splice site mutation |  |  |  |

Here we identified seven additional alleles in the *SVP* gene that enable the mutant plants to flower earlier despite experiencing a cold snap (**Fig. 5** and **Table 1**). Five new alleles were identified by whole-genome sequencing (*frosty1*, *frosty6*, *frosty7*, *frosty8*, and *frosty10*) while two new alleles were identified by sequencing the coding sequences of *SVP* in respective mutants (*frosty2* and *frosty4*). Three of the newly identified mutations (*frosty6*, *frosty7*, and *frosty10*) affect splice sites, while the other four induce point mutations (*frosty1*, *frosty2*, *frosty4*, and *frosty8*).

The *SVP* coding region of *frosty5*, *frosty11*, and *frosty12* were also sequenced but did not reveal any mutations. This suggests that another suppressor of flowering in response to a cold snap may be mutated in these mutants, or the mutation may affect parts of the *SVP* gene that were not sequenced, such as UTRs, promoter, or introns.

## Discussion

We have shown that the majority of the mutants we have identified here are impaired in floral repression in response to a cold snap and that the mutations we have identified by sequencing are all located in the *SVP* gene. The frosty genetic screen was conducted in the *eds1* mutant background. Although *eds1* mutants were previously found to have greater freezing tolerance (Chen et al., 2015), we observed no effect on their ability to flower earlier in response to a cold snap. On the contrary, we observed a mild late flowering phenotype in response to long cold days (**Fig. 1**), which may indicate that flower repression is enhanced in *eds1* mutant plants.

The observed changes in flowering time among the different *frosty* mutants were highly variable. However, the *frosty* mutants whose entire genomes were sequenced (*frosty1*, *frosty6*, *frosty7*, *frosty8*, *frosty10*) exhibited a consistent early flowering response in both cold long days and short-day conditions. Among these mutants, *frosty7* flowered similarly to wild type plants under long day conditions, while the other sequenced frosty mutants flowered uniformly early. As *frosty7* is a splice site mutation, these findings suggest that the respective splice isoform can substitute for a functional *SVP* gene and repress flowering under normal long day conditions, while the other mutants cannot. Interestingly, *frosty* mutants that had wild type *SVP* coding sequences (*frosty5*, *frosty11*, *frosty12*) also showed a more varied flowering response in both warm long and short days. The latter finding could indicate that other genes are affected in these mutants or that these mutants contain variations in regulatory sequences of the *SVP* gene that do not fully disrupt gene function. Mutations in *SVP* can present as semi-dominant, with heterozygotes showing a gradient in flowering time, ranging from early to wildtype flowering (Hartmann et al., 2000). We also observed this in the F2 generation of our *frosty* mutants, which, before knowing the identity of the gene that was mutated, puzzled us. For this reason, for whole genome sequencing we focused on *frosty* mutants that displayed the clearest 1:3 segregation pattern of the early flowering phenotype under cold long days. This pattern was observed in only five *frosty* mutants (*frosty1*, *frosty6*, *frosty7*, *frosty8*, *frosty10*) whilst we found it impossible to identify early flowering individuals in the *frosty2* and *frosty4* mutants and too few individuals were identified as early flowering in the rest of the mutants for us to confidently proceed with whole genome sequencing. Identifying causal mutations when the respective SNP is not homozygous or when the phenotype is not stable is substantially more difficult and bears the risk that the causal mutations remain elusive.

The *frosty9* mutant represents a special case, as mentioned above. It was initially identified as early flowering in cold-long days, but analysis of its progeny under the same conditions revealed no significant acceleration of flowering when considering the entire population. However, some individuals do flower more rapidly in cold long days, as well as long and short days at regular temperatures (see **Figs. 1 and 2**).

Homozygote second-site mutations were identified in *frosty1*, *frosty8* and *frosty10*. However, all mutations are in close proximity to *SVP*, and thus, they are most likely a result of genetic linkage. The second-site mutation in *frosty1* is in the gene for *DIACYLGLYCEROL KINASE 5* (*DGK5*; *AT2G20900*). DGK5 catalyzes the conversion of diacylglycerol to phosphatidic acid. Loss-of-function mutants have impaired resistance to pathogens and decreased freezing tolerance (Tan et al., 2018; Kalachova et al., 2022). However, no effect on flowering has been reported. Therefore, it is unlikely that a respective second-site mutation contributes to the flowering phenotype.

A second site mutation was found in *frosty8*, in a gene that encodes a cysteine/histidine-rich C1 domain family protein (*AT2G21850*). No loss of function phenotype has been described for this gene yet, and it appears to be primarily expressed in the root. *Frosty10* also carried a second mutation site in the gene for *MITOCHONDRIAL EDITING FACTOR 21* (*MEF21*; *AT2G20540*). MEF21 belongs to the family of pentatricopeptide repeat (PPR) proteins involved in mitochondrial mRNA editing. Due to genetic redundancy, the *mef21* loss-of-function mutant exhibits no discernible phenotypic differences from wild-type plants (Takenaka et al., 2010). Therefore, it seems that the observed flowering phenotypes in *frosty1*, *frosty8*, and *frosty10* are solely the result of mutations in the *SVP* gene and not due to second-site mutations.

Now that we have established that mutations in *SVP* are causal for the promotion of flowering in response to a cold snap, we investigated the nature of the mutations we identified. Three out of the five mutations identified in this study affect splice sites, while one results in a premature stop codon (*frosty1*, R96X) and another, *frosty8*, is a missense mutation that converts Alanine at position 32 to Valine (A32V). This mutation is noteworthy because it was previously identified as a QTL in the Japanese accession Fuk (Méndez-Vigo et al., 2013) and the same SNP was found to help a local accession of Arabidopsis to adapt to the Yangtze River basin (Zou et al., 2017). Genetic studies using an accession from Sichuan, China identified the exact same SNP and revealed additional roles for SVP in drought adaptation (Guo et al., 2023). It is worth noting that Alanine 32 is highly conserved in most Arabidopsis MADS-box domain transcription factors. Yet the amino acid sequence of AGAMOUS-LIKE (AGL) proteins AGL48, AGL80, AGL96 and AT5G27810 reveals a Valine at position 32. This supports the notion that the A32V mutation in SVP does not result in a loss-of-function mutation. Instead, it alters the DNA-binding ability, preventing the *SVP* variant in *frosty1* mutants and the Southeast Asian accessions from repressing their respective targets.

The E56K mutation was detected in *frosty4*, affecting an amino acid at the border of the MADS domain that is encoded at the end of exon 1. It is important to note that amino acid E56 is not highly conserved in other MADS-box proteins. Additionally, one of the annotated splice isoforms of *SVP* (*AT2G22540.2*) does not contain it. Due to the potent early flowering phenotype of *frosty4*, the latter observation may indicate that *SVP* encodes protein isoforms that are impaired in floral repression. These impaired proteoforms may act as decoys and sequester other protein interactors, such as FLM, allowing plants to fine-tune the degree of floral repression.

In addition to the MADS domain, the SVP protein also possesses a K domain that acts as an oligomerization module, allowing SVP to engage in homo- and heteromeric complexes (Lai et al., 2019). Ectopic expression of the K-domain alone can sequester MADS-box transcription factors into non-productive heteromers and thereby act in a dominant-negative fashion on their activity (Song and Chen, 2018). Our mutagenesis approach did not identify any SNPs affecting the K-domain.

However, splice site mutations were identified that affect the K-domain. In *frosty6*, the splicing defect resulted in a premature termination, leading to the loss of one-third of the K-domain. Meanwhile, *frosty10* had a sequence insertion in the K-domain as a consequence of intron retention. The K-domain is similar in structure to a three-helix leucine zipper. Therefore, removing parts or inserting any sequence into it will significantly impact the folding of the domain, which is essential for protein-protein interactions. It is possible that no SNPs were detected in the K-domain because only a few residues are necessary for the physical contacts between the helices of the leucine zipper. Mutations affecting residues outside of these may be more easily absorbed and not significantly affect protein function.

In sum, this work demonstrates the central role of SVP in cold snap-induced floral repression and identifies novel genetic variants, some of which affect the *SVP* splicing patterns. Our study provides significant insight into the molecular processes that regulate temperature-sensitive flowering in Arabidopsis, while also highlighting the dynamic relationship between *SVP* and environmental signals.

## Acknowledgements

We acknowledge funding through NovoCrops Centre (Novo Nordisk Foundation project number 2019OC53580 to S. W.), the Independent Research Fund Denmark (0136-00015B and 0135-00014B to S. W.) and the Novo Nordisk Foundation (NNF18OC0034226, NNF20OC0061440 and NNF23OC0084195 to S. W.).

